# Sense-and-respond payload delivery using a novel antigen-inducible promoter improves suboptimal CAR-T activation

**DOI:** 10.1101/2021.04.02.438280

**Authors:** Tingxi Guo, Dacheng Ma, Timothy K. Lu

## Abstract

Chimeric antigen receptor (CAR)-T cell therapies demonstrate the clinical potential of lymphocytes engineered with synthetic properties. However, CAR-T cells are ineffective in most solid tumors, partly due to inadequate activation of the infused lymphocytes at the site of malignancy. To selectively enhance anti-tumor efficacy without exacerbating off-target toxicities, CAR-T cells can be engineered to preferentially deliver immunostimulatory payloads in tumors. Here, we report a novel antigen-inducible promoter and single-vector sense-and-respond circuit for conditional payload expression in primary human T cells. In therapeutic T cell models, the novel NR4A-based promoter induced higher transgene expression than the conventional NFAT-based promoter under weakly immunogenic conditions, where payload expression is most needed. Minimal activity was detected from the inducible promoters in the absence of antigen and after withdrawal of stimulation. As a functional proof-of-concept, we used the NR4A-based promoter to express cytokines in an anti-mesothelin CAR-T model with suboptimal stimulation, and observed improved proliferation compared to T cells engineered with the conventional NFAT promoter or CAR alone. Our single-vector circuit achieves CAR-directed payload expression under weakly immunogenic conditions and could enable the next generation of cell therapies with enhanced anti-tumor efficacy.

---

Genetically programming cell functions with synthetic components holds promise for a variety of clinical applications. A notable example is the adoptive transfer of T lymphocytes engineered with a chimeric antigen receptor (CAR) to treat cancer^1,2^. However, the consistent clinical benefit of these therapies has been largely limited to hematological malignancies. Most carcinomas remain non-responsive to CAR-T cells because the suppressive tumor microenvironment and variable antigen density prevent adequate activation of lymphocytes^3, 4, 5^ Amplifying the suboptimal responses of therapeutic T cells without exacerbating immune-mediated toxicity is a major unmet need for the treatment of solid tumors.

Beyond antigen receptors, adoptively transferred anti-tumor T cells can also be engineered to produce immunostimulatory payloads^6^. This strategy to augment immune responses can enhance the therapeutic properties of the infused T cells and reinvigorate endogenous immune cells. In preclinical models, T cells engineered to secrete common gamma chain cytokines IL-2, IL-7, IL-15, and IL-21^7^; inflammatory cytokines IL-12^8^, IL-18^9^, and IL-23^10^; or other protein-based therapeutics^11^, ^12^ have demonstrated superior tumor control compared to non-producers. The continuous secretion of stimulatory payloads, however, may counteract their beneficial effects. In one case, human T cells engineered to constitutively produce IL-15 resulted in the transformation of transductants in an IL-15 receptor-dependent manner^13^. Constitutive production of potent cytokines such as IL-2 or IL-18 also caused toxicities in pre-clinical CAR-T models^7, 9^. These observations highlight the need to tightly control recombinant payload production and, ideally, restrict it to the tumor site to maximize its clinical benefit and prevent unwanted side-effects^14^.

A synthetic biology approach could leverage the antigen receptor signaling in engineered lymphocytes to specify where the stimulatory payloads should be produced. A sensitive, antigen-inducible promoter with low background activity could localize payload delivery. The conventional approach for antigen-dependent transgene expression has been to use an NFAT-based promoter^15^ encoding an NFAT/AP1 response element derived from the human IL-2 enhancer^16^. This NFAT promoter was tested in the clinic to drive inducible expression of IL-12 and was transduced to *ex vivo* expanded tumor-infiltrating lymphocytes^17^. Toxicities were still observed after infusion, possibly because of non-localized production of IL-12 by T cells with unknown antigen specificities. Subsequent pre-clinical developments have focused on combining the antigen-inducible NFAT promoter with a recombinant receptor^9, 14, 18^ to better control the input signal for conditional payload expression. Despite its broad use, the standard NFAT promoter may not be the optimal choice for payload delivery.

Here, we identified a novel synthetic promoter based on an NR4A-binding motif that induced greater responses than the conventional NFAT promoter under weakly stimulatory conditions, which is when immune-enhancing molecules are most needed. We also describe a single-vector circuit that incorporates this synthetic promoter to achieve an automated payload response via CAR sensing of the cognate tumor antigen. The engineered T cells respond to targets in an antigen-dependent manner and conditionally express a transgene of choice upon antigen engagement. The inducible promoter and vector design described here could enable future generations of synthetic lymphocytes, with controllable input and output to safely enhance therapeutic responses.

## RESULTS

### A novel antigen-inducible promoter encoding an NR4A-binding motif

We previously generated a synthetic promoter library termed Synthetic Promoters with Enhanced Cell-State Specificity (SPECS), based on transcription factor (TF) binding motifs found in public databases. SPECS vectors were constructed by encoding repeated TF binding sites upstream of a minimal promoter derived from the adenoviral major late promoter (MLP) and mKate as the fluorescent reporter^19^. In the present study, to identify novel antigen receptor-inducible promoters from this library, we selected individual candidate promoters encoding binding sites for known TFs directly downstream of T cell receptor (TCR) signaling pathways (i.e., NFkB and MAPK targets)^16, 20, 21, 22, 23, 24, 25, 26^, or TFs upregulated upon TCR-induced activation^27, 28, 29, 30, 31, 32, 33, 34^. TF binding site sequences ranged from 77-126 base pairs (bp) (Supplementary Table 1). Individual promoter vectors were transduced into primary human T cells by lentivirus and stimulated with plate-bound CD3 agonist OKT3, or left untreated as a control (Supplementary Fig. 1A). Among the tested promoters, the one encoding an NR4A-binding motif induced the highest percentage of reporter positive cells compared to other candidates (Supplementary Fig. 1B and 1D) and was selected for characterization. An AP1-based promoter was also chosen for comparison, because the AP1 pathway is well-established in T cell activation^35^. As an internal positive control, CD137 upregulation^36^ was measured in all assays to ensure similar activation among experiments (Supplementary Fig. 1C).

Next, we compared the SPECS-derived NR4A and AP1 promoters with the conventional NFAT promoter for OKT3-inducible responses. To facilitate quantitative comparisons, we introduced a second downstream transcription module into the lentiviral vector. In this module, the constitutive EFS promoter drives expression of truncated CD271 (tCD271) to mark transduced cells. We also tested an additional synthetic minimal promoter (SMP)^37^ in combination with each of the three response elements (Fig. 1A). The SMP and a similar variant enabled robust inducible promoter activity in human cells^18, 38^. All of the promoter vectors transduced cells with comparable efficiency at ~60-80% (Supplementary Fig. 2A). As a negative control vector, we cloned the EFS-tCD271 module alone, without inducible promoters or mKate. All of the tested promoters responded similarly to the CD3 agonist among CD4 and CD8 subsets of primary human T cells after one day of stimulation (Fig. 1B). In certain donors or in a TF-dependent context, SMP performed better than MLP, although no consistent differences were observed (Fig. 1B and 1C). Thus, subsequent experiments were performed with SMP as the minimal promoter. No significant baseline activity in the absence of stimulation was observed with any of the inducible promoters (Fig. 1B). Using a set of vectors with only a minimal promoter sequence upstream of mKate, we did not detect enhancer-like activity from the constitutive EFS promoter at the steady-state or after activation, regardless of the choice of minimal promoter (Supplementary Fig. 3).

**Figure 1.**
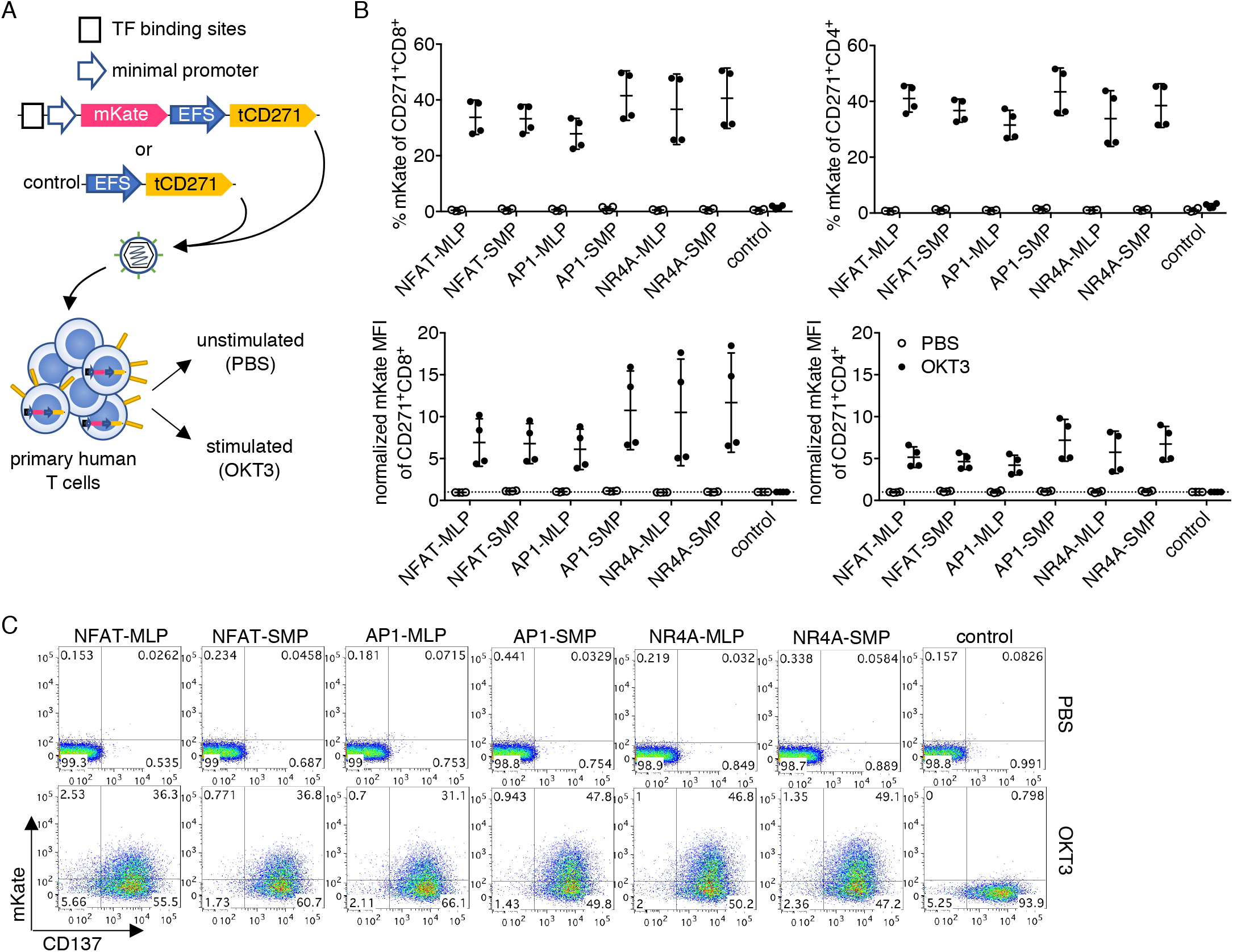
NFAT, AP1, and NR4A-based promoters are activated by anti-CD3 stimulation with minimal background. **a** Vector and experimental schematics. Response elements encoding NFAT, AP1, or NR4A-binding sites were cloned with either a core promoter from the adenovirus-derived major late promoter (MLP) or a synthetic minimal promoter (SMP). Lentiviral vectors were transduced into primary human T cells, and cells were treated with PBS or anti-CD3 clone OKT3. **b** Reporter fluorescence in transduced (CD271+) CD8+ or CD4+ cells was measured after 24 hours. **c** Representative flow plots gated on CD271+CD8+ T cells are shown. Lines and error bars denote mean ± standard deviation. n=4 from 2 independent donors tested in 2 technical replicates.

To investigate whether the TCR-inducible promoters could be activated by non-CD3 dependent mechanisms, we cultured the promoter-transduced cells in conditioned media derived from strongly activated T cells to mimic an inflammatory milieu (Supplementary Fig. 4A). The NFAT, AP1, and NR4A promoters were activated at similarly low levels (~10%) when the transduced cells were cultured in the conditioned media compared to normal media. Reporter activity induced by the conditioned media was substantially lower than CD3-induced responses (Supplementary Fig. 4B and 4C). Thus, we have identified a novel NR4A-based promoter with anti-CD3 inducible activity, and a single lentiviral vector system that permits stringent conditional gene expression alongside constitutive gene expression.

### Inducible promoters demonstrate reversible and repeatable activation

We next investigated the activity of the inducible promoters after withdrawal of antigen receptor stimulation. Using the vectors shown in Figure 1A, we observed that it took up to 5 days after removing the source of stimulation for the mKate fluorescence to dissipate (Supplementary Fig. 2B), suggesting high stability of the fluorescent protein. To measure the reversibility and repeatability of inducible promoter activation, we changed the reporter to a destabilized enhanced yellow fluorescent protein (dEYFP) encoding an additional PEST motif, which reduces the half-life of fluorescent proteins^39^. In this system, the reporter fluorescence is more closely coupled to promoter activity. After transducing the dEYFP vectors in human T cells, we stimulated the cells for one day with OKT3 as above, then transferred the cells to a fresh well without the agonist for three days of rest. This process was repeated three times (Fig. 2A). The EFS-tCD271 vector again served as a negative control. Across three sequential stimulations, the NFAT, AP1, and NR4A promoter activities consistently returned to baseline after three days of rest in CD8 T cells. In fact, the fluorescence intensity for all reporters was reduced by at least 50% after only one day (Fig. 2B-D).

**Figure 2.**
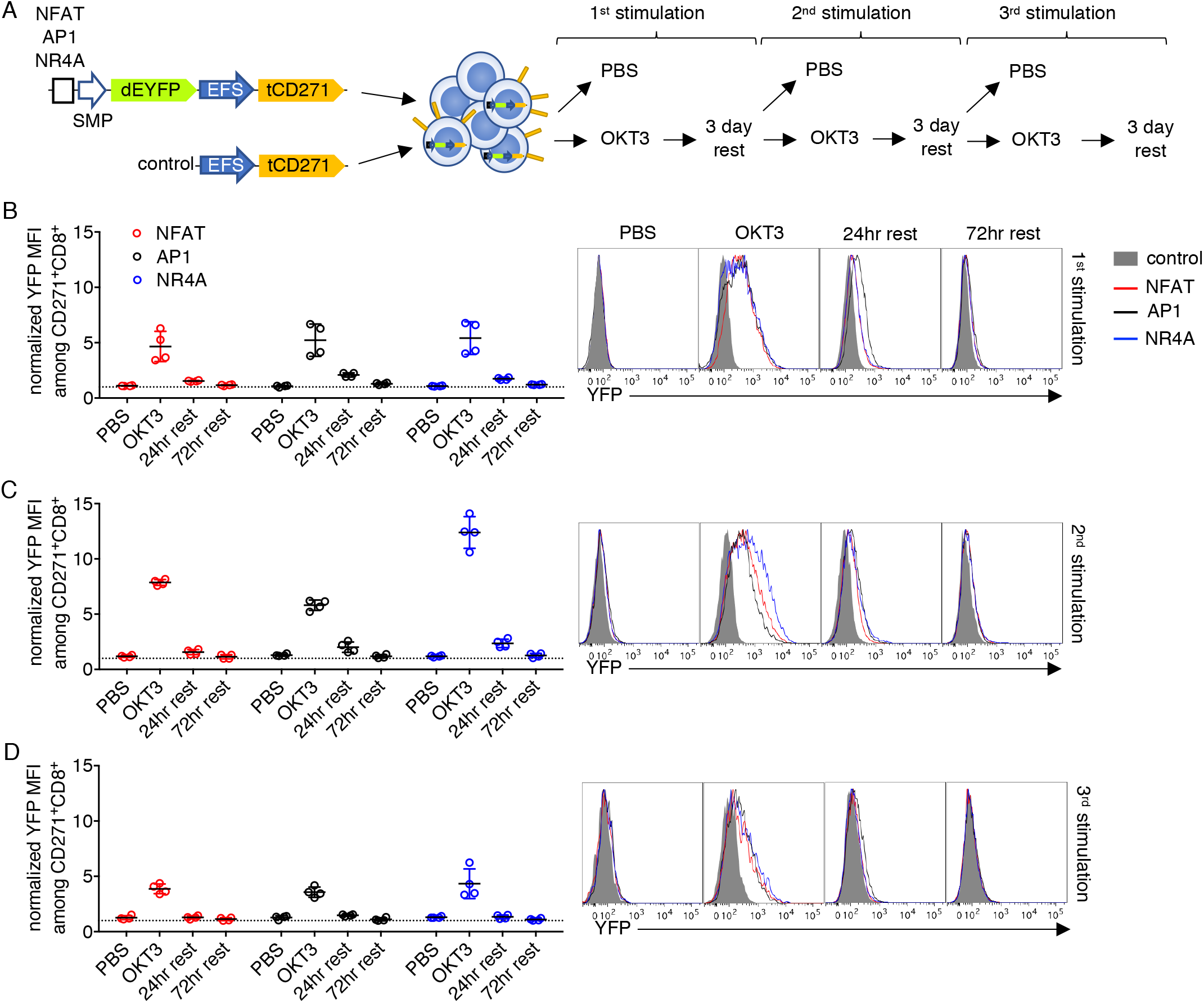
Inducible promoter responses are reversible and repeatable. **a** Vector and experimental schematics. Response elements encoding NFAT, AP1, or NR4A-binding sites with the synthetic minimal promoter (SMP) were used to drive destabilized yellow fluorescence protein (dEYFP). Lentiviral vectors were transduced into primary human T cells, and cells were treated with PBS or anti-CD3 clone OKT3 for 24 hours, then transferred to a fresh plate to rest for up to 3 days before repeating the process two more times. **b-d** Reporter fluorescence in CD271+CD8+ cells was measured after 24 hours of stimulation, then after 24 and 72 hours of rest, following the first (**a**), second (**b**), and third (**c**) round. Fluorescence intensity was normalized to that of control vector transductants. Representative histograms gated on CD271+CD8+ T cells are shown. Lines and error bars denote mean ± standard deviation. n=4 from 2 independent donors tested in 2 technical replicates.

Interestingly, normalized responses were moderately higher after the second stimulation (Fig. 2B, 2C, and Supplementary Fig. 5A), akin to a recall response in adaptive lymphocytes. The lower responses observed after the third stimulation (Fig. 2D and Supplementary Fig. 5A) were likely the result of activation-induced cell death. Reversible responses were also observed in CD4 T cells with at least one round of stimulation (Supplementary Fig. 5B). Repeated OKT3 stimulation biased the outgrowth of CD4-cells and decreased the overall viability of most samples (Supplementary Fig. 5C); thus, the promoter responses in the CD4 subset after multiple rounds of activation could not be reliably measured. Throughout the course of the experiment, the proportion of CD271+ cells did not change significantly (Supplementary Fig. 5D), indicating that repeated activation of the promoters was well tolerated and did not cause a growth disadvantage.

### NR4A promoter induces higher responses than NFATand AP1 in weakly immunogenic, therapeutically relevant models

Although OKT3 is a potent activator of T cells, it is not representative of therapeutically relevant receptor-antigen interactions. To characterize the inducible promoter responses in more appropriate models, we first selected a CAR based on the humanized single-chain variable fragment (scFv) M5, targeting the widely expressed mesothelin tumor antigen^40^. The M5 CAR, which encodes 41BB and CD3z signaling domains (M5-BBz), is currently being tested in clinical trials for treating a variety of solid tumors (NCT03054298, NCT03323944).

Mesothelin-targeting CAR-T strategies have yet to yield consistent objective responses^41^ and, therefore, could benefit from the addition of inducible payloads. To investigate the inducible promoter activity in a CAR setting, the M5-BBz CAR and tCD271, separated by a porcine 2A (P2A) sequence, were encoded downstream of the EFS promoter for constitutive expression. In these constructs, inducible promoters that drive the expression of mKate as a reporter were cloned upstream of the CAR. tCD271 with the receptor alone served as a control vector (Fig. 3A). T cells were transduced with the vectors and stimulated with HEK293T (no mesothelin), A549 (low mesothelin), or OVCAR8 (high mesothelin) target cells^42, 43^.

**Figure 3.**
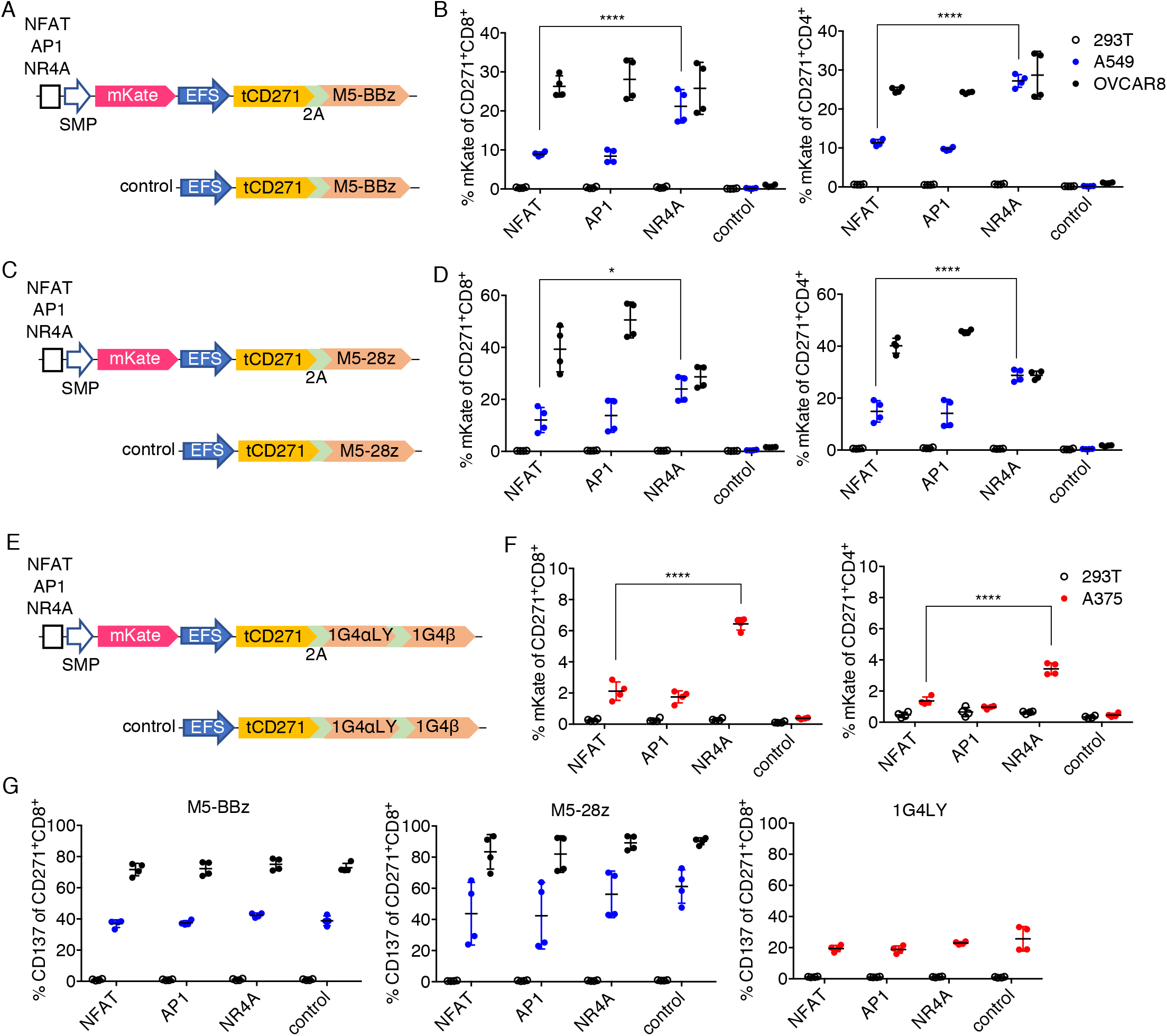
NR4A-based promoter induces higher responses than NFAT or AP1 in weakly stimulatory TCR/CAR-T models. **a,c,e** Schematics for CAR or TCR and inducible module encoded within a single vector. **b,d,f** Primary human T cells transduced with the vectors shown on the left of respective graphs were co-cultured with the indicated target cells, and mKate fluorescence was measured after 24 hours for CD271+CD8+ or CD4+ subsets. **g** CD137 expression on CD271+CD8+ T cells from the same experiments shown in panels **b**, **d**, and **f.** Lines and error bars denote mean ± standard deviation. *P<0.05, ****P<0.0001 by two-way ANOVA adjusted for all possible comparisons using Tukey’s test. n=4 from 2 independent donors tested in 2 technical replicates.

Among the transduced CD8 and CD4 T cells, we observed no mKate fluorescence when effector cells were cultured with HEK293T, again demonstrating minimal antigen-independent promoter activity (Fig. 3B). Similar levels of reporter expression were observed with the strong OVCAR8 stimulation. Notably, the NR4A promoter induced significantly more mKate expression than the NFAT and AP1 promoters when cultured with the weakly stimulatory A549 targets (Fig. 3B and Supplementary Fig. 6A). OVCAR8 was indeed more immunogenic than A549 in the M5 model based on CD137 upregulation (Fig. 3G and Supplementary Fig. 6A). Doubling the number of NFAT binding sites only marginally improved the response, which was still lower than that of NR4A in the M5-BBz/A549 system (Supplementary Fig. 7). A similar trend was observed when the promoters were tested with a CD28-based M5 CAR (M5-28z, Fig. 3C): the NR4A promoter responded at higher levels than the standard NFAT in response to A549 stimulation (Fig. 3D and Supplementary Fig. 6A). In the M5-28z model, AP1 demonstrated higher activity than NR4A in response to OVCAR8 (Fig. 3D). Inducible promoter vectors for both CARs were transduced at ~50-70% efficiency (Supplementary Fig. 6B).

Next, we constructed a similar set of vectors using the affinity-matured HLA-A2/NYESO1-specific 1G4 TCR^44^ as the model antigen receptor. 1G4 TCR has demonstrated clinical efficacy in treating melanoma and synovial sarcoma^45, 46^ The two P2A sequences between tCD271, TCRα, and TCRβ genes were codon-modified to avoid repetition in the viral genome (Fig. 3E). TCR-T cells were stimulated with HEK293T or A375. Both of these cell lines express HLA-A2 but only A375 expresses the cognate antigen^44, 47^. In the TCR model, NR4A also responded with consistently higher reporter positivity than NFAT or AP1 after coculture with A375, although the overall responses were lower than those seen with the CAR models. The promoters induced higher responses in CD8 T cells than in CD4 cells (Fig. 3F), which was expected since the 1G4 TCR is HLA class I restricted. The lower levels of CD137 upregulation in the TCR model were consistent with the weaker promoter activity (Fig. 3G), which may have been caused by the insufficient formation of the correct TCRα/β pairing (see Discussion). The TCR constructs were ~400bp larger than the CAR vectors and were transduced less efficiently (Supplementary Fig. 6B). Nevertheless, across all of the receptor models tested in our study, we observed significantly higher responses with the NR4A-based promoter compared to the responses observed with the standard NFAT promoter under poorly stimulatory conditions — precisely the context where payload expression is needed.

### Recombinant cytokines expressed under the NR4A promoter amplify weak anti-tumor responses

As a proof-of-concept, we generated inducible constructs to conditionally express IL-2 and IL-21 in the clinically relevant M5-BBz model. The mKate reporter gene of the M5-BBz CAR vectors (Fig. 3A) was replaced with recombinant IL-2 and IL-21, separated by a P2A sequence (Fig. 4A). Constitutive expression of either cytokine alone has been reported to enhance the proliferation of CD19 CAR-T cells^7^. Although both of these common gamma chain cytokines can be produced endogenously by activated human T cells, cytokine production is poor when the cells are suboptimally stimulated at low antigen density^48, 49, 50^. Therefore, we reasoned that the NR4A-based synthetic promoter system could supplement these beneficial cytokines under conditions that preclude endogenous production.

**Figure 4.**
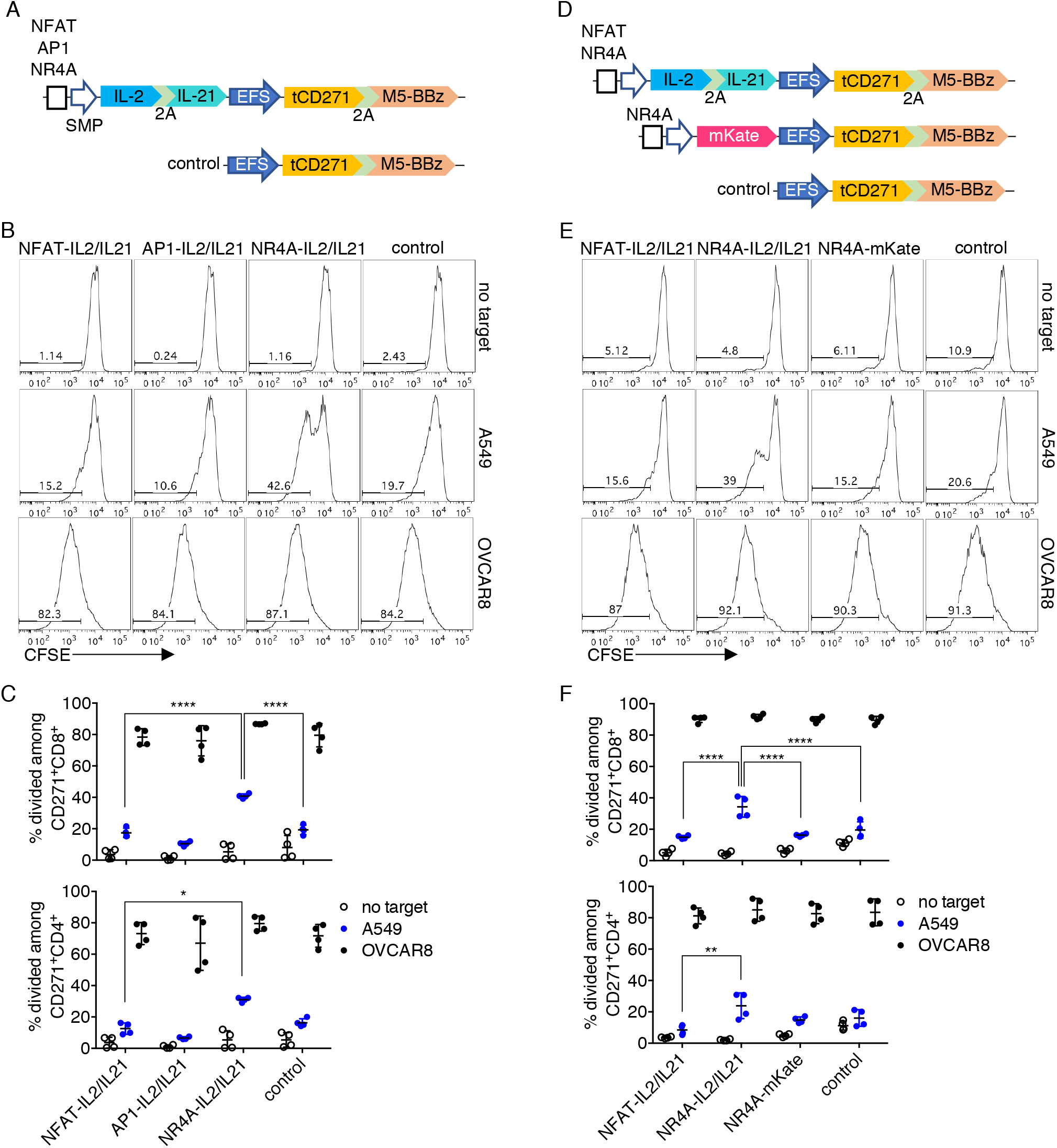
Cytokines delivered by the NR4A-based promoter amplify weak antigen-specific responses. **a** Schematics for vectors encoding M5-BBz CAR with or without IL-2 and IL-21 as inducible payloads. **b** Primary human T cells transduced with the vectors shown in **a** were labeled with CFSE and co-cultured with the indicated target cells. Dye dilution was measured after 4 days of co-culture. Representative histograms gated on CD271+CD8+ T cells are shown. **c** Quantification of CFSE dilution in CD271+CD8+ or CD4+ subsets. **d** Schematics for vectors encoding IL-2 and IL-21 as inducible payloads or mKate as a control. **e** Experiment was performed as described in **b**. Representative histograms gated on CD271+CD8+ T cells are shown. **f** Quantification of CFSE dilution in CD271+CD8+ or CD4+ subsets. Lines and error bars denote mean ± standard deviation. *P<0.05, **P<0.01, ****P<0.0001 by two-way ANOVA adjusted for all possible comparisons using Tukey’s test. n=4 from 2 independent donors tested in 2 technical replicates.

In a proliferation assay without cytokine supplementation in the media, CAR-T cells transduced with the control M5-BBz or inducible IL2/IL21 vectors demonstrated low levels of proliferation in the absence of stimulation. In contrast, the majority of the cells divided after co-culture with the immunogenic OVCAR8 cells. When stimulated with the weakly immunogenic A549 targets, more of the NR4A-IL2/IL21 transductants proliferated compared to cells engineered with other vectors. The improvement in proliferation was more pronounced for the CD8 than CD4 subset (Fig. 4B and 4C), consistent with a past study showing that IL-2 improves the proliferation of suboptimally stimulated CD8 but not of CD4 murine T cells^51^. Based on additional experiments in which the NR4A promoter induced expression either of cytokines or mKate as a control (Fig. 4D), we determined that the enhanced proliferation in response to A549 was payload-dependent, at least in CD8+ CAR-T cells (Fig. 4E and 4F). Transduction efficiencies of the inducible cytokine constructs were similar (Supplementary Fig. 8 and 9C).

Consistent with the above data, more of the CAR-T cells encoding the NR4A-IL2/IL21 module produced IL-2 when stimulated with A549, compared to cells transduced with other inducible modules or the control vector (Supplementary Fig. 9A and 9B). Moreover, in A549 co-cultures, proliferation of NFAT-IL2/IL21 CAR-T cells was not increased compared to control CAR-T cells (Fig. 4C and 4F), in line with the low responses observed with the NFAT promoter in Figure 3B. Across these experiments, inducible expression of the cytokines did not significantly enhance proliferation compared to CAR alone when cells were cultured with OVCAR8 (Fig. 4C and 4F). In summary, these data demonstrate a proof-of-concept that payloads delivered via the single-vector system using the NR4A promoter can augment suboptimal CAR-T responses.

## DISCUSSION

In this study, we identified a novel antigen-inducible transcriptional response element in human T cells based on a TF binding site of the NR4A family. Notably, the NR4A-based promoter outperformed the conventional NFAT-based promoter under poorly stimulatory conditions. Given that at least NR4A1 expression is partly dependent on calcium and MAPK signaling^52^, it is interesting that our NR4A-based promoter was more responsive than NFAT and AP1-based promoters. The NR4A family of TFs could simply be more potent transcriptional activators in primary human T cells. Alternatively, the choice of TF binding motif or vector design used here may promote transcription for NR4A more than for NFAT or AP1. Future molecular and biochemical studies are required to elucidate the mechanism underlying the differential activity between respective promoters at various signaling intensities. NFAT and AP1-based synthetic promoters may also be disadvantageous because they sequester critical TFs away from endogenous response elements that are needed to potentiate activation, especially with weak signals. Meanwhile, NR4A TFs have been implicated as negative regulators of T cell activation^53, 54^ and thus the cognate synthetic promoter would be unlikely to interfere with endogenous activation pathways.

TCR-induced activation of the NR4A pathway has been characterized previously^31, 52, 55^. This pathway has been used to monitor TCR signaling by knocking-in a reporter at the NR4A1 locus^55, 56, 57^. In theory, payload transgenes could also be knocked-in at the NR4A1 site to achieve inducible expression, and improvements in site-specific integration technologies for primary lymphocytes^58, 59, 60^ will facilitate the practical implementation of this approach. With the knock-in method, however, transcriptional output will be dictated by endogenous elements, which lacks the flexibility afforded by a synthetic promoter system that can be tuned for a variety of applications. Our promoter platform similarly leverages the NR4A pathway; instead of relying on endogenous response elements to drive NR4A1 transcription, a short sequence encoding an NR4A-binding motif is used to drive conditional gene expression, which is readily implemented using a standard lentiviral vector. Compared to other systems in which the inducible module is transduced via a vector separate from the antigen receptor^14^, our single-vector platform simplifies the manufacturing process and avoids the heterogeneity in the final therapeutic product generated by multi-vector transduction.

The EFS promoter was used here to constitutively express the recombinant antigen receptor. At ~200bp, it is one of the most compact constitutive promoters; the small size is advantageous for incorporation into the large vectors designed in this study. Other compact constitutive promoters, such as MND, tend to possess enhancer-like activity and can influence the transcriptional activity of neighboring promoters^61, 62, 63, 64^. Such cross-distance activity would interfere with the upstream promoter for achieving stringent inducible transcription. Although the EFS promoter is weaker than MND^65^, we did not detect enhancer-like properties, consistent with published data^64^. However, stronger constitutive promoters may be needed to practically implement this system with recombinant TCRs, which must compete with endogenous TCR hemichains to form antigen-specific surface receptors^66^. The low level of response observed in our 1G4 model is likely caused by insufficient expression of TCRα/β chains by the EFS promoter. Efforts to engineer potent and compact constitutive promoters without enhancer-like attributes, or inducible promoters that are resistant to distal enhancers, are needed.

In our proof-of-concept experiment, the inducible expression of common gamma chain cytokines by the NR4A promoter improved proliferation of otherwise poorly responsive CAR-T cells. Activation at a lower immunogenic threshold is a critical feature of the NR4A antigen-inducible promoter that could widen the therapeutic index for a wide range of molecular therapeutics compared to systemic or constitutive delivery. Some examples of applications include site-specific production of: bispecific engagers to trigger bystander lymphocyte responses; chemokines to promote infiltration of immune effectors; or cell-intrinsic regulators (e.g., TFs) to conditionally re-program engineered cells in an autonomous manner. These and other applications can be explored in future pre-clinical studies. In conclusion, the platform described in this study could empower a variety of synthetic biology approaches to overcome current challenges in adoptive cell immunotherapies.

## METHODS

### Cell culture

HEK293T, A375, A549, and Jurkat.E6 cell lines were obtained from the American Type Culture Collection (ATCC, Manassas, VA). OVCAR8 was a gift from Dr. Sangeeta N. Bhatia (Massachusetts Institute of Technology, Cambridge, MA). HEK293T, A375, A549, and OVCAR8 cell lines were cultured in DMEM (Thermo Fisher Scientific, Waltham, MA; catalog #10569010). Primary human T cells and the Jurkat.E6 cell line were cultured in RPMI-1640 (Thermo Fisher Scientific; catalog #11875119). All media were supplemented with 10% fetal bovine serum (Corning; catalog #35-01 0-CV) and 1% penicillin/streptomycin (Thermo Fisher Scientific; catalog #15140122).

### Generation of lentiviral vectors

Truncated CD271 (tCD271)-CAR or TCR, EYFP destabilized with a PEST motif (dEYFP), and IL-2/IL-21 fragments were synthesized as gBlocks by Integrated DNA Technologies (Coralville, IA). The M5 scFv sequence was derived from the patent WO2015/090230. Sequences of 28z and BBz signaling domains, and the affinity-matured 1G4 TCR were as previously described^44, 67^. Except NFAT, all TF binding site and mKate sequences were derived from SPECS plasmids (Addgene #127842). The NFAT response element was subcloned from the pSIRV-NFAT-eGFP plasmid^68^ (a gift from Peter Steinberger, Addgene plasmid # 118031). The EYFP sequence was derived from Addgene plasmid #51791 and the PEST sequence was derived from Addgene plasmid #69072. The EFS promoter was subcloned from the lentiCRISPRv2 plasmid (a gift from Feng Zhang, Addgene plasmid # 52961). The EFS promoter and tCD271-CAR/TCR fragments were first assembled into a lentiviral backbone vector (derived from pFUGW in-house) in the reverse orientation of the long-terminal repeats. Inducible promoter and reporter or payload genes were then inserted upstream of the EFS promoter. Inserted sequences were confirmed by Sanger sequencing (GENEWIZ, South Plainfield, NJ). Each set of NFAT, AP1, or NR4A vectors only differed at the TF binding sequence.

### Lentivirus production

Lentivirus was generated by transfecting HEK293T cells of less than 10 passages and grown to ~80% confluency in T25 flasks, with 1.5μg of pMD2.G (a gift from Didier Trono, Addgene plasmid # 12259), 3.5μg of psPAX2 (a gift from Didier Trono, Addgene plasmid # 12260), and 5μg of respective transfer plasmid using the TransIT-2020 reagent (MirusBio, Japan). Media was changed to fresh complete DMEM 16-24 hours post-transfection, and lentivirus was collected after another 24 hours to be used immediately or stored at −80°C. Virus was titered by infecting Jurkat.E6 at limiting dilutions.

### Lentiviral transduction of primary human T cells

Peripheral blood mononuclear cells (PBMCs) were isolated by density gradient centrifugation from apheresis products of healthy donors (Brigham and Women’s Hospital Crimson Core, Boston, MA). Primary human T cells were purified from PBMCs by Pan T Cell Isolation Kit (Miltenyi Biotec, Germany). Purified T cells were stimulated with anti-CD3/CD28 Dynabeads (Thermo Fisher Scientific) at a T cell:bead ratio of 1:2. After 24 hours, Dynabeads were removed and stimulated T cells were seeded on Retronectin (Takara Bio, Japan) coated nontissue culture treated plate with virus, and centrifuged at 1200xg, 32°C for 30 min. One time infection was carried out for the smaller vectors shown in Figures 1 and 2 at a multiplicity of infection (MOI) of 5-10. Larger vectors shown in Figure 3 and 4 were infected at a MOI of 10-20, spread out over 2 days of daily infection. During stimulation and infection, T cells were supplemented daily with 100U/ml of IL-2 and 10ng/ml of IL-15 (NCI Preclinical Repository, Frederick, MD). After infection, T cells were maintained by supplementing with 100U/ml of IL-2 and 10ng/ml of IL-15 every 3 days. T cells were expanded for another 5-6 days after the last infection prior to use in experiments.

### Flow cytometry

The following monoclonal antibodies (BioLegend, San Diego, CA) were used in this study: anti-human CD3 (clone UCHT1), anti-human CD4 (clone RPA-T4), anti-human CD8 (clone RPA-T8), anti-human CD271 (clone ME20.4), anti-human CD69 (clone FN50), anti-human CD137 (clone 4B4-1), and anti-human IL-2 (clone MQ1-17H12). Surface staining was carried out at 4°C for 15 min with a master mix of antibodies. For intracellular staining of IL-2, cells were fixed and permeabilized after surface staining using the Cytofix/Cytoperm kit (BD Biosciences). Stained cells were analyzed with a FACSCantoII, FACSCelesta, or LSRFortessa flow cytometer (BD Biosciences). Data analysis was performed with FlowJo. All data shown were gated on singlets and live cells, determined by Aqua fixable dye (Thermo Fisher Scientific) for mKate-expressing and intracellular experiments, or 7-aminoactinomycin D (BioLegend) for all other experiments.

### T cell stimulation assays

For plate-bound stimulations, non-tissue culture treated plates were coated with 2μg/ml anti-CD3 clone OKT3 (NCI Preclinical Repository) with or without 2μg/ml anti-CD28 clone CD28.2 (BioLegend) by incubating at 4°C overnight. The same volume of PBS (Thermo Fisher Scientific) was used as a control treatment. Antibody or PBS solution was removed and T cells were seeded to the treated wells and cultured for 24 hours. Cells were analyzed poststimulation or transferred to fresh tissue-culture treated wells to rest. For cell-based co-culture stimulations, T cells were mixed with HEK293T, A375, A549, or OVCAR8 targets at an effector:target (ET) ratio of 3:1 and cultured for 24 hours to measure reporter fluorescence. For intracellular IL-2 detection, CAR-T cells were cultured with targets at a 3:1 ET ratio for 18 hours, followed by treatment with 1500x diluted monesin (BD Biosciences) for 6 hours. To measure proliferation, T cells were washed with PBS and labeled with 1μM of carboxyfluorescein succinimidyl ester (CFSE, Thermo Fisher Scientific) by incubating them at 37°C for 5 min. Labeled cells were washed with complete media and co-cultured with A549 or OVCAR8 target cells at an ET ratio of 10:1. CFSE dilution was assessed after 4 days of coculture.

### Statistics

Comparisons between more than two groups were performed by two-way analysis of variance (ANOVA) with Tukey’s multiple comparisons test. Differences were considered significant at an adjusted P value of less than 0.05. All statistical analyses were performed using GraphPad Prism 6. Error bars denote one standard deviation.

## ACKNOWLEDGEMENTS

This work was supported in part by the Department of Defense (W81XWH-17-1-0159, W81XWH-16-1-0565) and National Institutes of Health (5-R33-AI121669-04). T.G. was supported by a Postdoctoral Fellowship from the Natural Sciences and Engineering Research Council of Canada. We thank the flow cytometry core facility at the Koch Institute for Integrative Cancer Research at Massachusetts Institute of Technology. We thank Karen Pepper for editing the manuscript.

## AUTHOR CONTRIBUTIONS

T.G., D.M., and T.K.L. designed the experiments, analyzed the data, and wrote the manuscript. T.G. and D.M. performed the experiments. T.K.L. supervised the study.

---

**Supplementary Figure 1.**
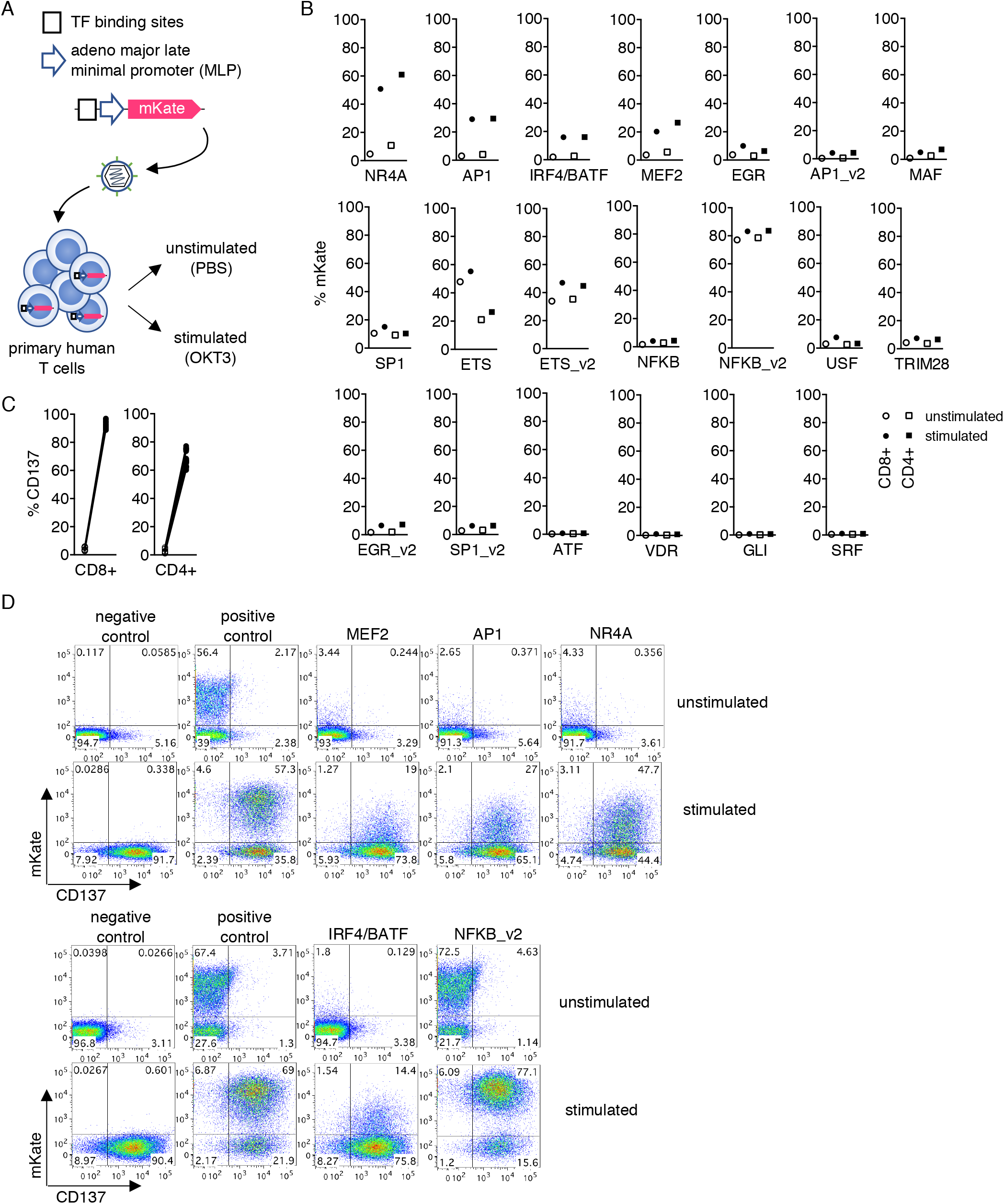
Screening for novel TCR-inducible response elements. **a** Schematics for vector design from the SPECS library and experimental setup. **b** Primary human T cells transduced with a panel of vectors were treated with PBS (unstimulated, open symbols) or OKT3 (stimulated, filled symbols) for 24 hours, and mKate fluorescence in the CD8+ (circles) or CD4+ (squares) subset was quantified. **c** CD137 expression after PBS (open) or OKT3 (filled) treatment. Measurements are matched by the same vector. **d** Raw flow plots gated on CD8+ T cells for several notable promoters are shown. The positive control vector encodes the constitutive UBC promoter to express mKate, whereas the negative control vector lacks mKate.

**Supplementary Figure 2.**
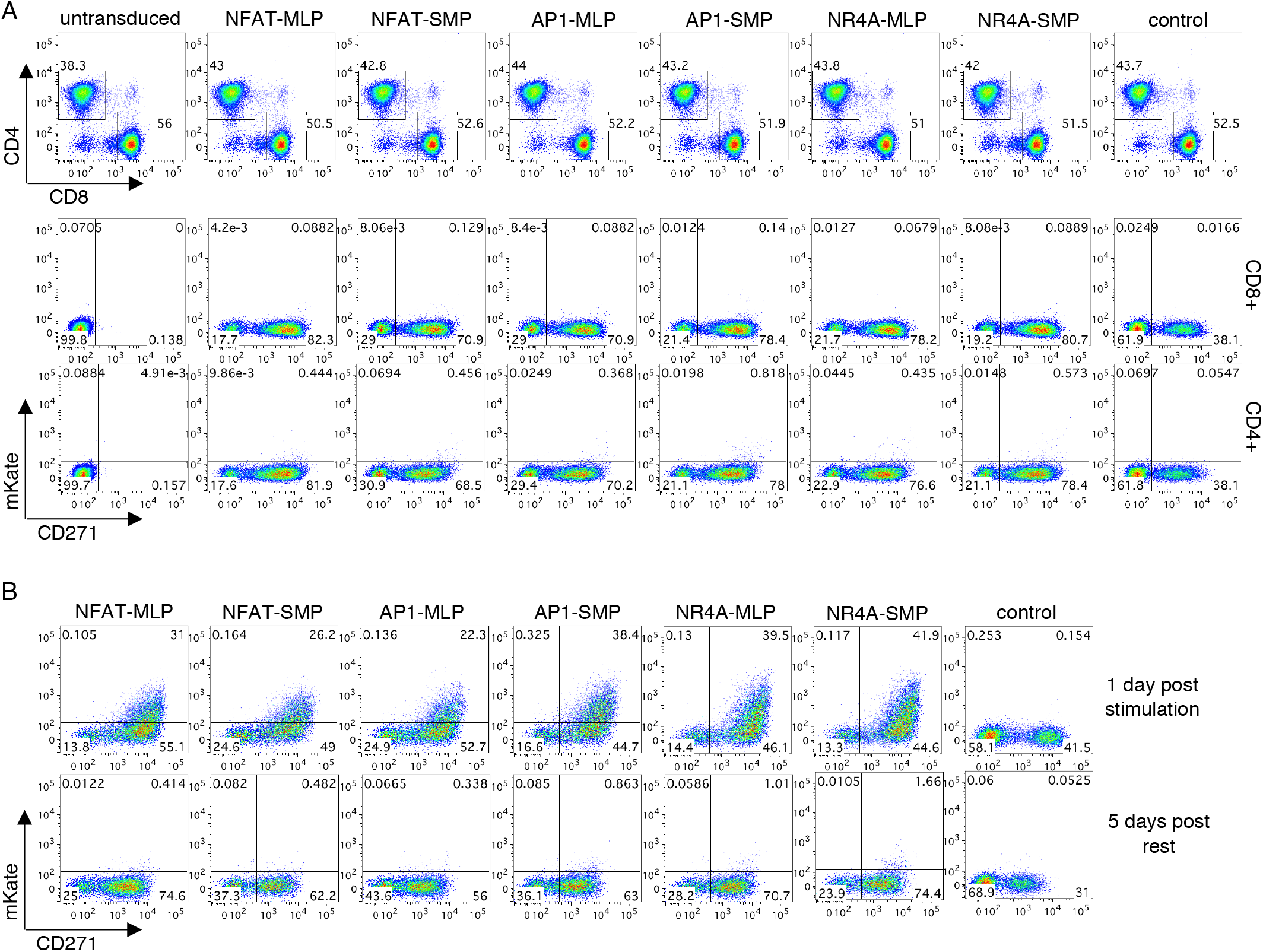
Gating strategy, transduction efficiency, and mKate stability. **a** CD4+ or CD8+ T cells are gated as shown on the top row. Transduction efficiency measured by CD271 positivity is shown on the bottom rows. **b** Fluorescence of the mKate reporter after one day of OKT3 stimulation and after 5 days of rest upon transfer to fresh wells is shown for representative CD8+ cells.

**Supplementary Figure 3.**
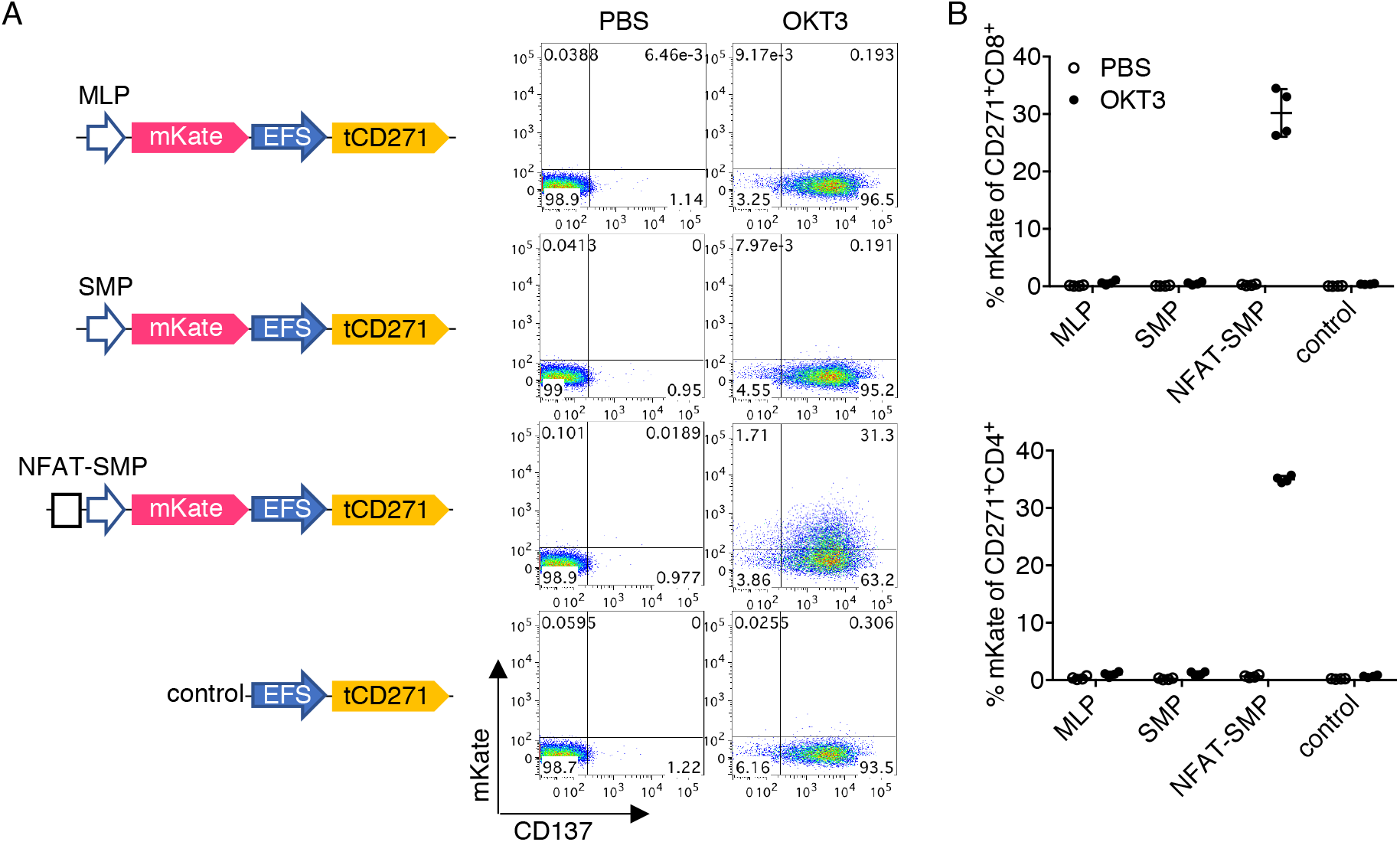
The EFS constitutive promoter do not possess detectable enhancer-like activity. **a** Primary human T cells transduced with the indicated vector shown on the left were treated with PBS or OKT3. CD137 and mKate upregulation was measured after 24 hours. Representative flow plots gated on CD271+CD8+ cells are shown on the right. **b** Quantification of data shown in panel **a**. Lines and error bars denote mean ± standard deviation. n=4 from 2 independent donors tested in 2 technical replicates.

**Supplementary Figure 4.**
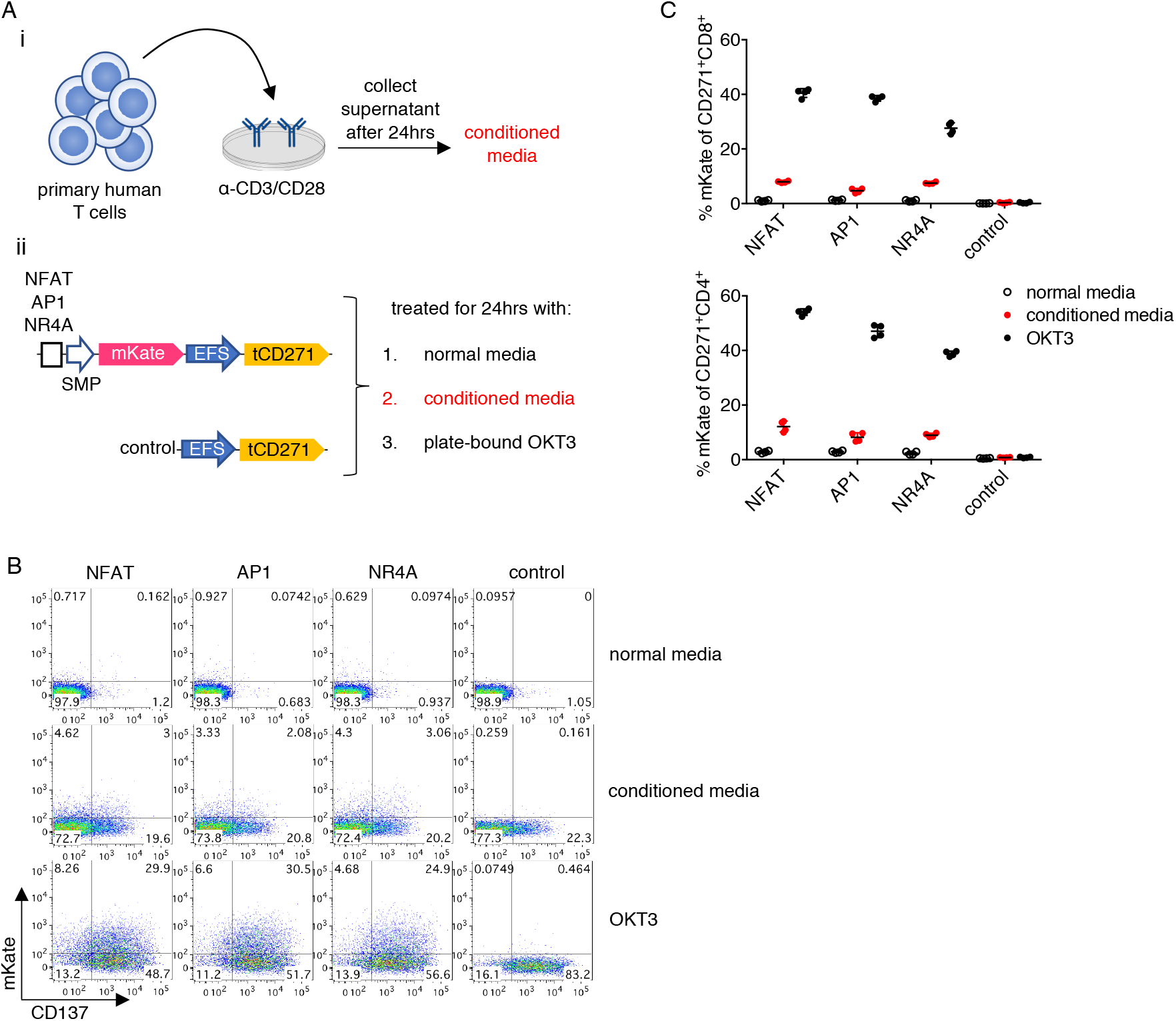
TCR-inducible promoters are weakly activated by an inflammatory milieu. **a** Experimental design. i) Conditioned media was generated by collecting the supernatant from expanded primary human T cells restimulated with plate-bound anti-CD3/CD28 monoclonal antibodies. ii) Promoter or control transduced T cells were cultured with normal media, conditioned media, or platebound OKT3. **b** Representative flow plots gated on CD271+CD8+ cells are shown. **c** Quantification of data shown in panel **b**. Lines and error bars denote mean ± standard deviation. n=4 from 2 independent donors tested in 2 technical replicates.

**Supplementary Figure 5.**
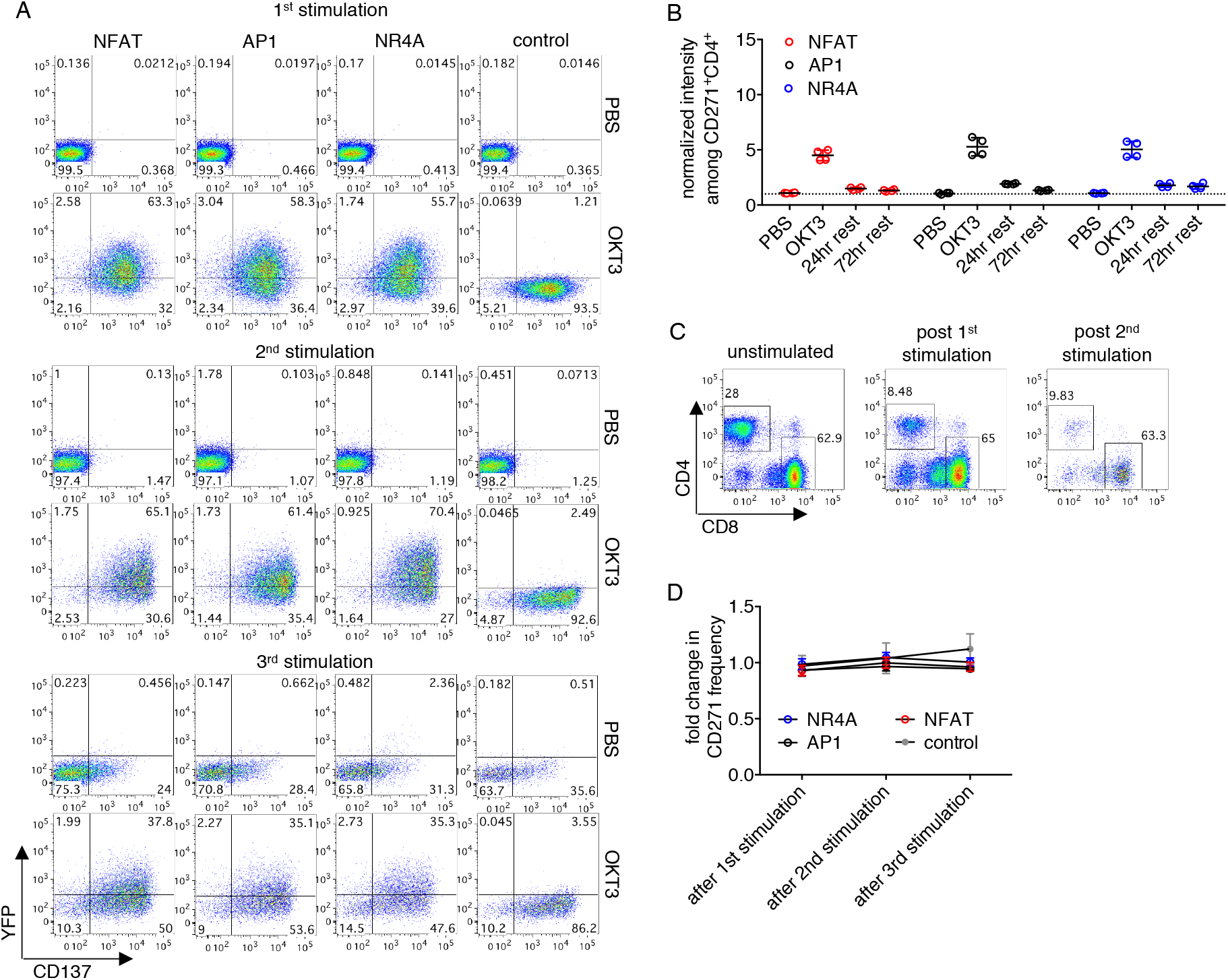
Inducible promoters can be repeatedly stimulated without causing cellular toxicity. **a** Representative flow plots gated on CD271+CD8+ cells for the PBS and OKT3 conditions in Figure 2B-D are shown. **b** Reversible promoter responses among CD271+CD4+ T cells were measured, as described in the caption of Figure 2, after the first round of stimulation. **c** Representative frequency of CD4+ and CD8+ subsets in culture after repeated stimulation. Gated on live cells. Due to the activation-induced cell death and biased outgrowth of CD4-cells, CD4+ cells could not be reliably analyzed after the first round of stimulation. **d** Change in percent of transduced (CD271+) cells was tracked through the three rounds of stimulation and remained constant. Frequency was normalized to that at the start of the experiment (prior to first stimulation). Lines and error bars denote mean ± standard deviation. n=4 from 2 independent donors tested in 2 technical replicates.

**Supplementary Figure 6.**
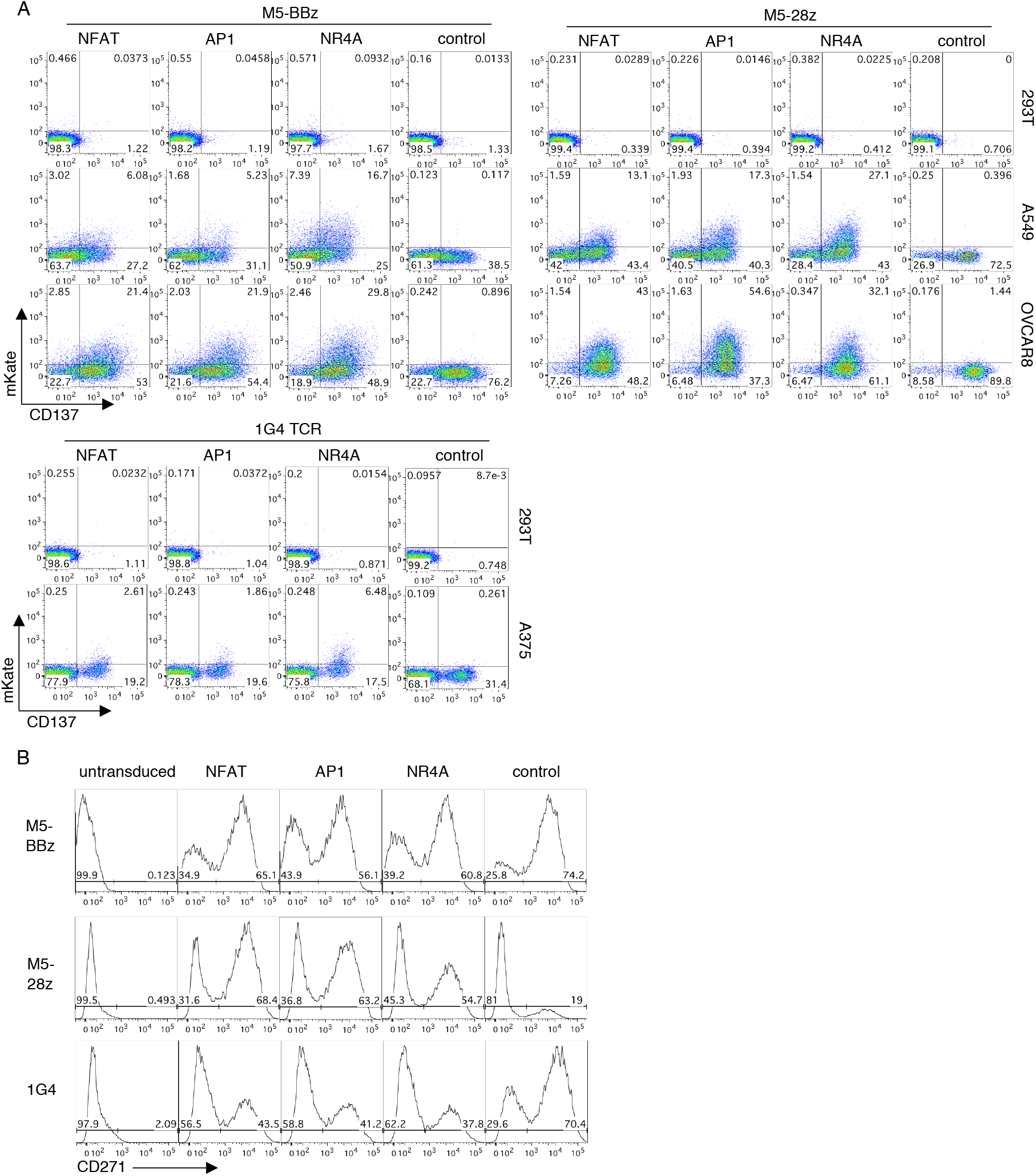
Inducible promoter activities in TCR/CAR-T models. **a** Representative flow plots gated on CD271+CD8+ cells for data in Figure 3 are shown. **b** Representative transduction efficiencies of the TCR/CAR vectors. Gated on total live cells. Note that the control M5-28z transduction was unexpectedly low. However, this does not affect the interpretation of the data, as the control M5-28z vector mainly serves as a gating control for mKate encoding vectors.

**Supplementary Figure 7.**
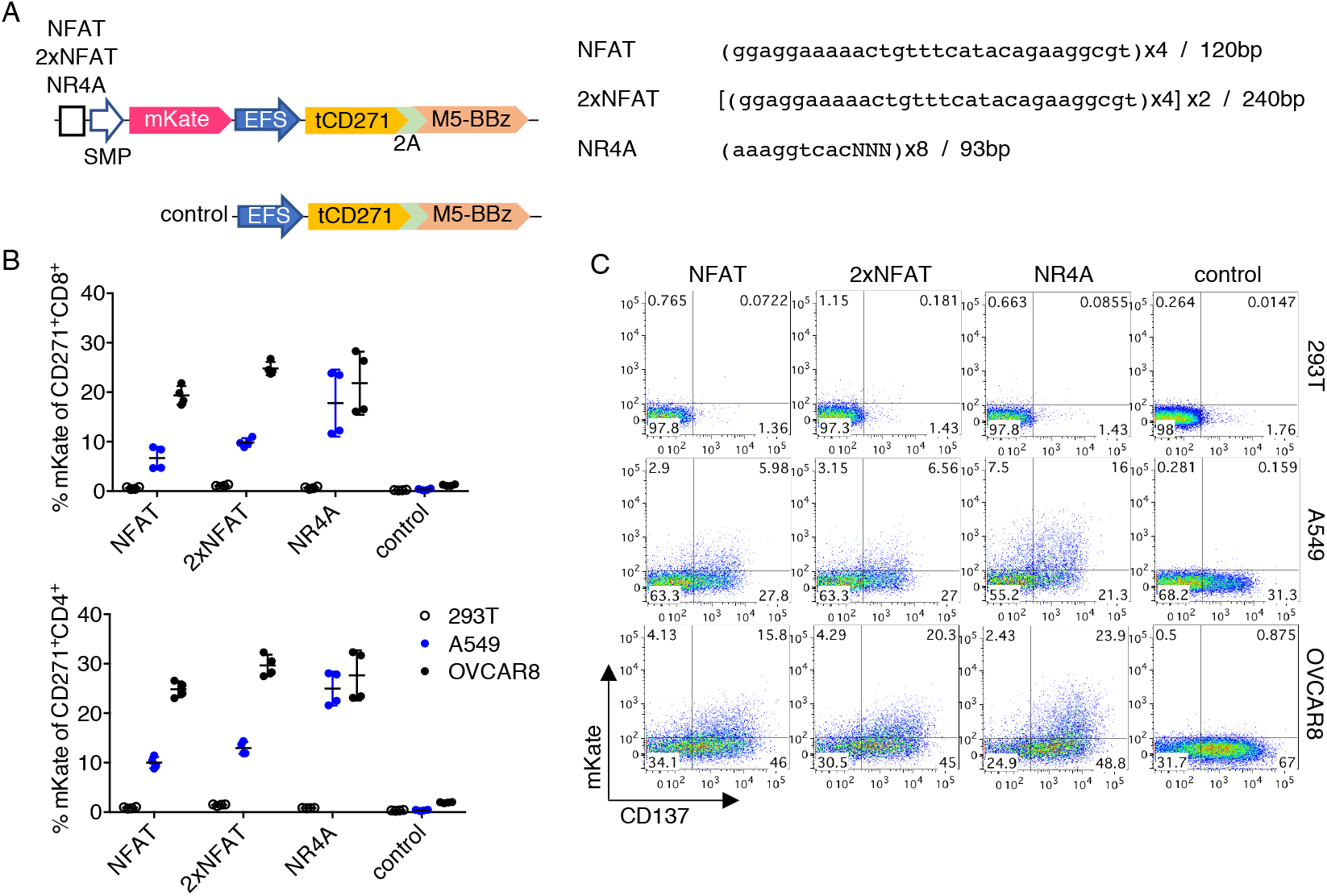
Doubling the number of NFAT binding sites only marginally improves response. **a** Left, vector schematics. Right, promoter sequences and designs. Two copies of the original NFAT promoter with 4 binding sites were cloned to generate 2xNFAT. The NR4A promoter was derived from the SPECS library, where transcription factor binding motifs were spaced by three random nucleotides. **b** Quantification of reporter responses among CD271+CD8+ or CD4+ subsets. **c** Representative flow plots gated on CD271+CD8+ cells for data shown in **b**. Lines and error bars denote mean ± standard deviation. n=4 from 2 independent donors tested in 2 technical replicates.

**Supplementary Figure 8.**
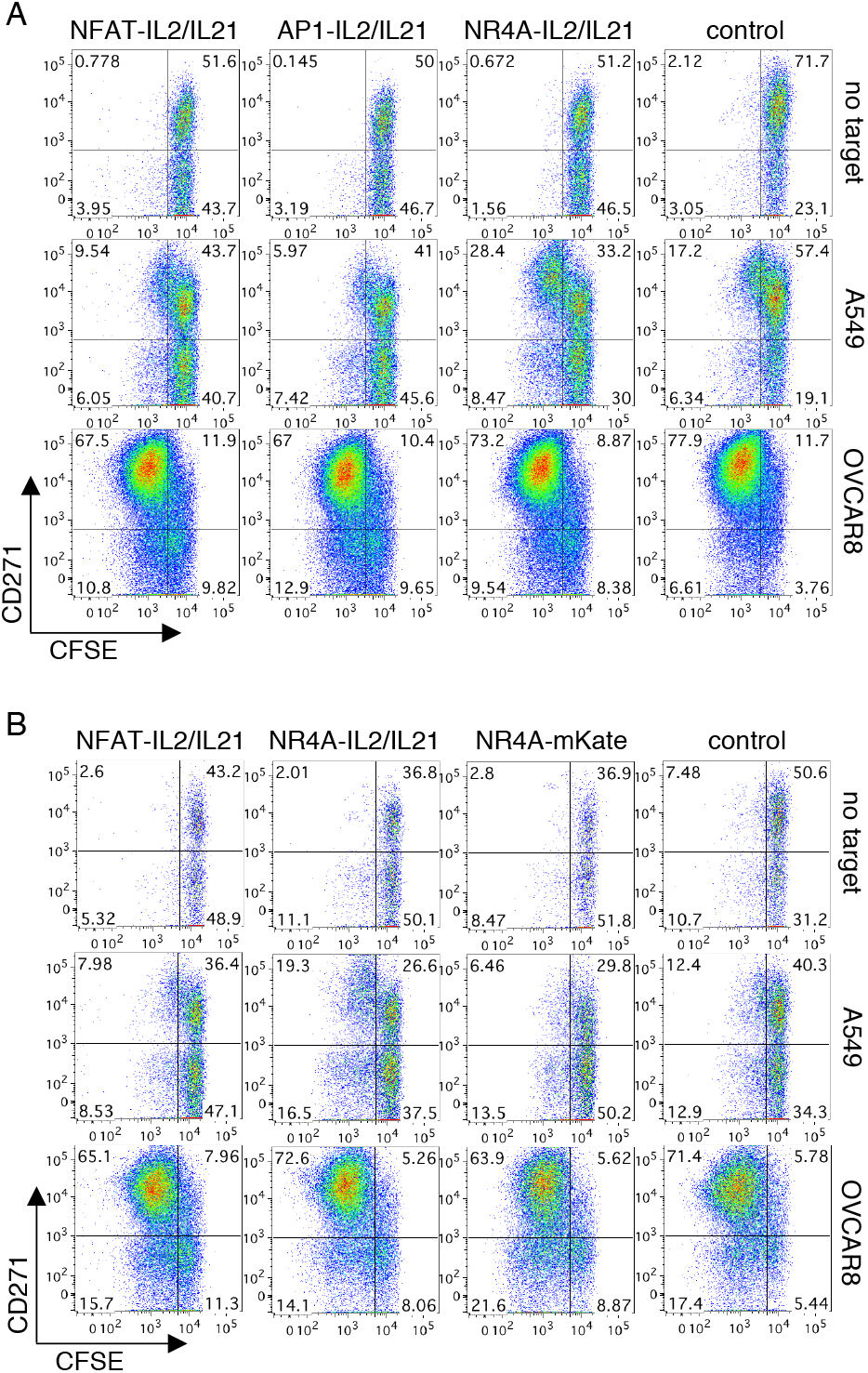
Inducible expression of cytokines augments weak antigen-specific response in CAR-T cells. **a** Representative flow plots gated on CD8+ cells for data in Figure 4A-C are shown. **b** Representative flow plots gated on CD8+ cells for data in Figure 4D-F are shown.

**Supplementary Figure 9.**
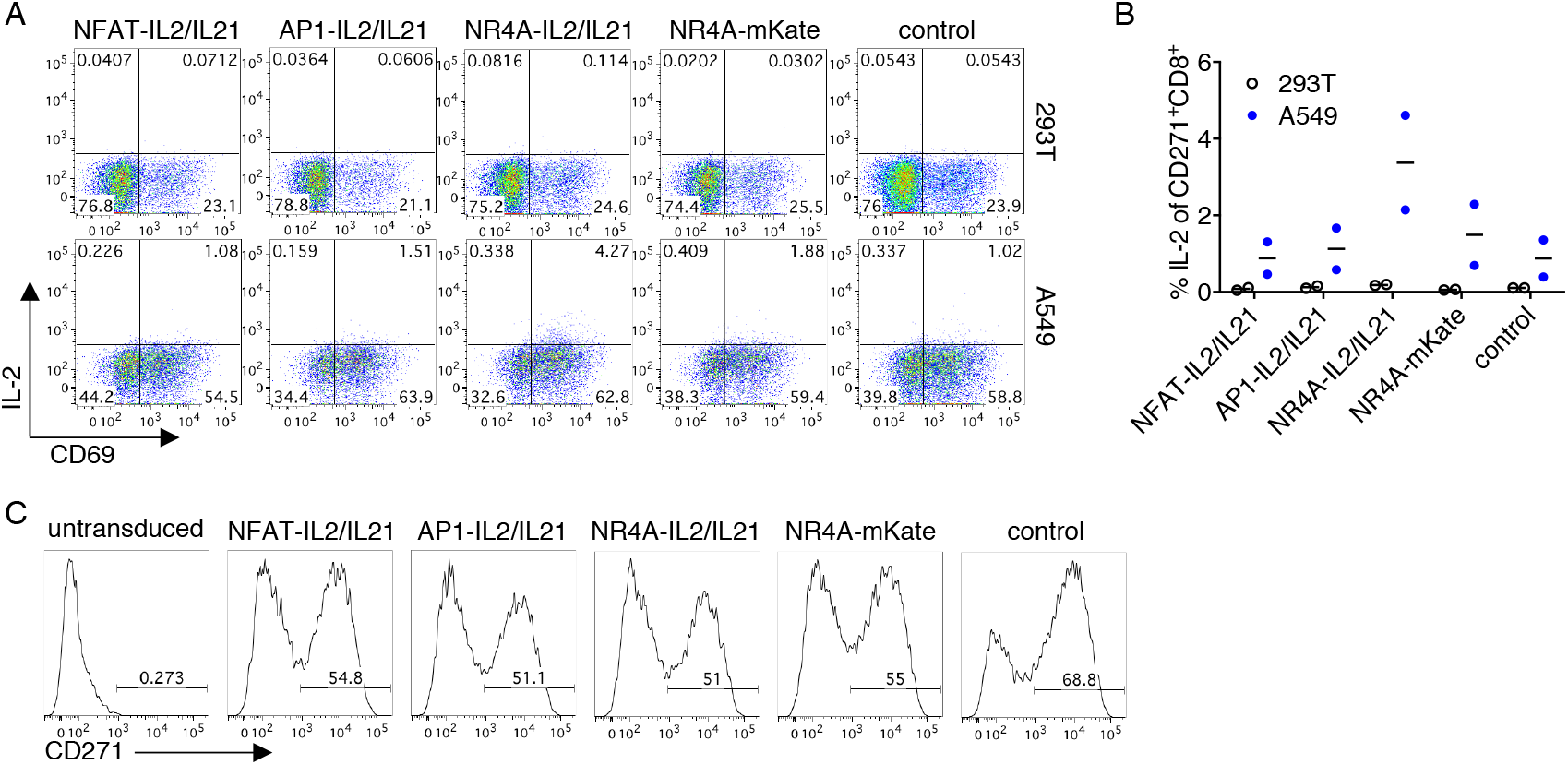
Analyses of inducible payload expression in stimulated CAR-T cells. **a** CAR-T cells transduced with inducible IL-2/IL-21 or control constructs were stimulated with 293T or A549 for 18 hours. Monesin was then added, and cells were cultured for another 6 hours before staining. Representative flow plots gated on CD271+CD8+ cells are shown. **b** Quantification for data shown in **a** across two independent donors. Lines denote means. **c** Representative transduction efficiencies of vectors used in this experiment and Figure 4.

**Supplementary Table 1.**
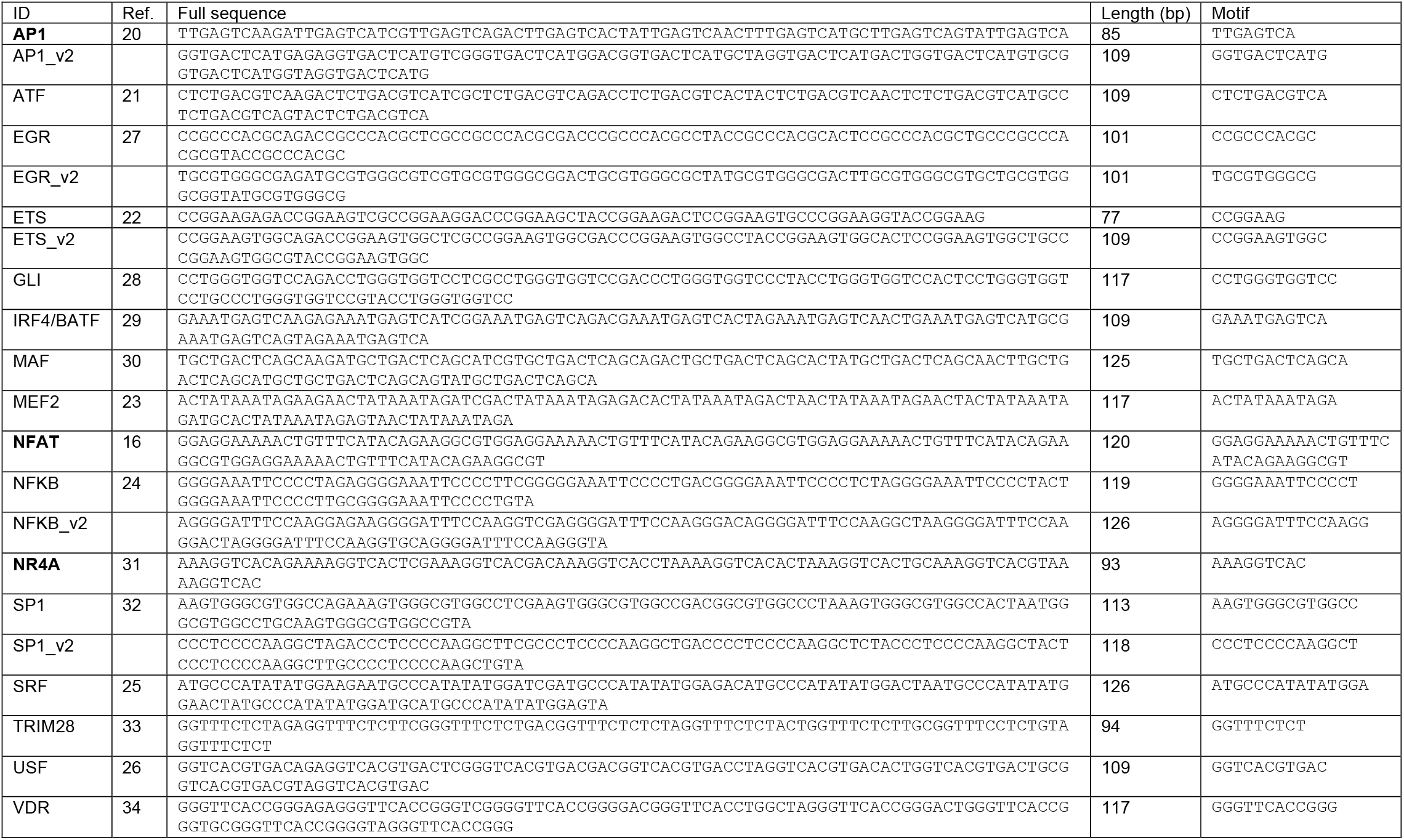
Transcription factor (TF) binding site sequences. Sequences and length for the promoters studied in Supplementary Figure 1 and the conventional NFAT promoter are shown. Bolded promoters were studied in the main figures. References for association of respective TF pathway with TCR signaling are listed.

